# Plumage colour structures global patterns of mixed-species bird flocks

**DOI:** 10.64898/2026.08.25.746914

**Authors:** Kanika Aggarwal, Imran Samad, Maria Thaker, Kartik Shanker

## Abstract

Mixed-species groups (MSGs) pose a particular challenge for our understanding of sociality in animals. Though MSGs are widespread social assemblages that form to enhance foraging success and reduce predation risk of participants, the role of traits in mediating grouping has received less attention. In particular, the role of body colour has not been tested quantitatively, despite the fact that visual similarity can reduce individual predation risk. Here, we examine whether plumage colour structures mixed-species bird flocks (MSFs) at a global scale. Using data spanning four continents, we developed a new metric that quantifies colour similarity among flock participants and compared observed flocks to null assemblages constructed from all flocking species at each site. We further examined whether MSF participants represented a colour subset of the available colours in the regional species pool. We found striking evidence that birds in MSFs were more similar in colour than expected by chance across all sites, indicating that plumage colour is a non-random structuring trait that shapes assembly of flocks globally. The strength and prevalence of colour structuring varied across geographies, but not flock size. Within communities, MSF participants differed systematically in colour composition from the regional species pool, occupying a restricted region of colour space dominated by yellow and brown plumage. Thus, plumage colour affects MSFs influencing both overall flock participation as well as species co-occurrence within flocks. Our findings illustrate the importance of visual traits in structuring interspecific social systems, by highlighting that birds of a feather do indeed flock together.

**Significance Statement:** Mixed-species bird flocks are among the most widespread examples of heterospecific social behaviour, yet the role of visual traits in shaping these associations remains poorly understood. Using a global dataset on bird colour and flock data spanning 52 sites across four continents, we developed a metric to test whether bird species participating in mixed-species flocks are more similar in plumage colour than expected by chance. We found consistent evidence that flocking species are significantly more colour-similar than random assemblages and occupy restricted regions of colour space from the local species pool. These patterns were consistent across geographical sites. Our findings suggest that visual similarity may contribute to flock cohesion or predator avoidance, highlighting the role of phenotypic traits in structuring ecological communities and heterospecific groups.

## Introduction

Grouping behaviour occurs in many taxa and confers multiple benefits, and yet group participation can be quite selective. Animal groups may comprise single species (conspecific groups), or may involve members of different species, known as mixed-species groups (hereafter, MSGs) or heterospecific groups. Such mixed-species groups have been observed across the animal kingdom from butterflies to fish, birds, and mammals. Mixed-species groups of forest birds or mixed-species flocks (hereafter, MSFs) are typically an interactive community of largely insectivorous birds which move and forage together to gain enhanced access to resources and protection from predation^2^. These roving groups with at least two species participants are found in terrestrial habitats globally^1^ and vary widely in size, stability, and strength of interspecific associations^3,4,5^.

At a global scale, MSF participants have been found to be more similar than expected by chance, particularly in phenotypic traits such as morphology (e.g., body size) and behavioural traits (e.g., foraging behaviour), suggesting that birds may largely join flocks for supplementary anti-predator benefits^2,5^. Participation in flocks can also reduce predation risk via positional effects, dilution of individual attack probability, increased collective vigilance, and reduced predator targeting efficiency^5, 6,7^. Since MSFs are dynamic, visually-coordinated groups that rely on rapid information transfer, collective movement, and mutual detection, traits related to visual signalling and detectability are likely to play a key role in mediating flock cohesion and assembly^2,5,8^. Plumage colour, therefore, represents a promising yet largely unexplored axis along which MSF assembly may be structured.

Birds are among the most visually striking organisms on Earth, exhibiting exceptional diversity in plumage colour that spans nearly the entire avian-visible spectrum^9^. Avian colouration is evolutionarily persistent across lineages^9,10^, and plays a central role in avian communication and signal evolution^10,11,12^, with ecological consequences for detectability and predator–prey interactions^13^. Comparative analyses further show that plumage colouration is shaped by the combined effects of ecological selection, sexual selection, and life-history strategies^10^. Moreover, the presence of specific colours across avian lineages can be predicted by evolutionary history and ecological context, including habitat type, light environment, and signalling background^9^. Thus, avian colouration reflects ecological adaptation and evolutionary constraints, representing a key functional trait.

If functional traits influence who associates with whom in ecological communities, then plumage colour should be expected to play a role in MSF formation^14,15^. Traits such as body size, foraging strata, and vocal behaviour are known to structure participation and roles within MSFs, shaping patterns of information flow, vigilance, and predator detection^1,3,16,17^. However, visual traits—despite being central to avian perception, communication, and predator–prey interactions—have rarely been examined as potential drivers of flock assembly. As a result, it remains unknown whether plumage colour functions as a community-level trait filter in MSFs, shaping both within-flock assembly and the subset of species that participate in flocks from the regional species pool^14,18^.

Here, we test whether plumage colour functions as a structuring visual trait in MSFs at a global scale (see Fig. 1 for a summary). Using data from 52 sites spanning four continents, we combine null-model simulations, colour similarity metrics, and community-level comparisons to address three related questions. First, we ask whether species participating in MSFs are more colour-similar within flocks than expected by chance, indicating non-random, colour-structured assembly at the level of individual flocks. Second, we assess whether the strength of colour filtering varies with flock size and across biogeographic regions, testing whether colour-based structuring is context dependent. Finally, we examine whether the colours of MSF participants represent a random subset of the regional species pool, or occupy a biased or restricted region of colour space relative to the community. By integrating within-flock and community-level analyses, our study evaluates whether plumage colour acts as a multi-scale trait filter shaping both the internal structure and broader composition of MSFs. In doing so, we extend trait-based approaches to social community assembly and provide a general framework for understanding how visual traits influence interspecific social systems.

**Figure 1:**
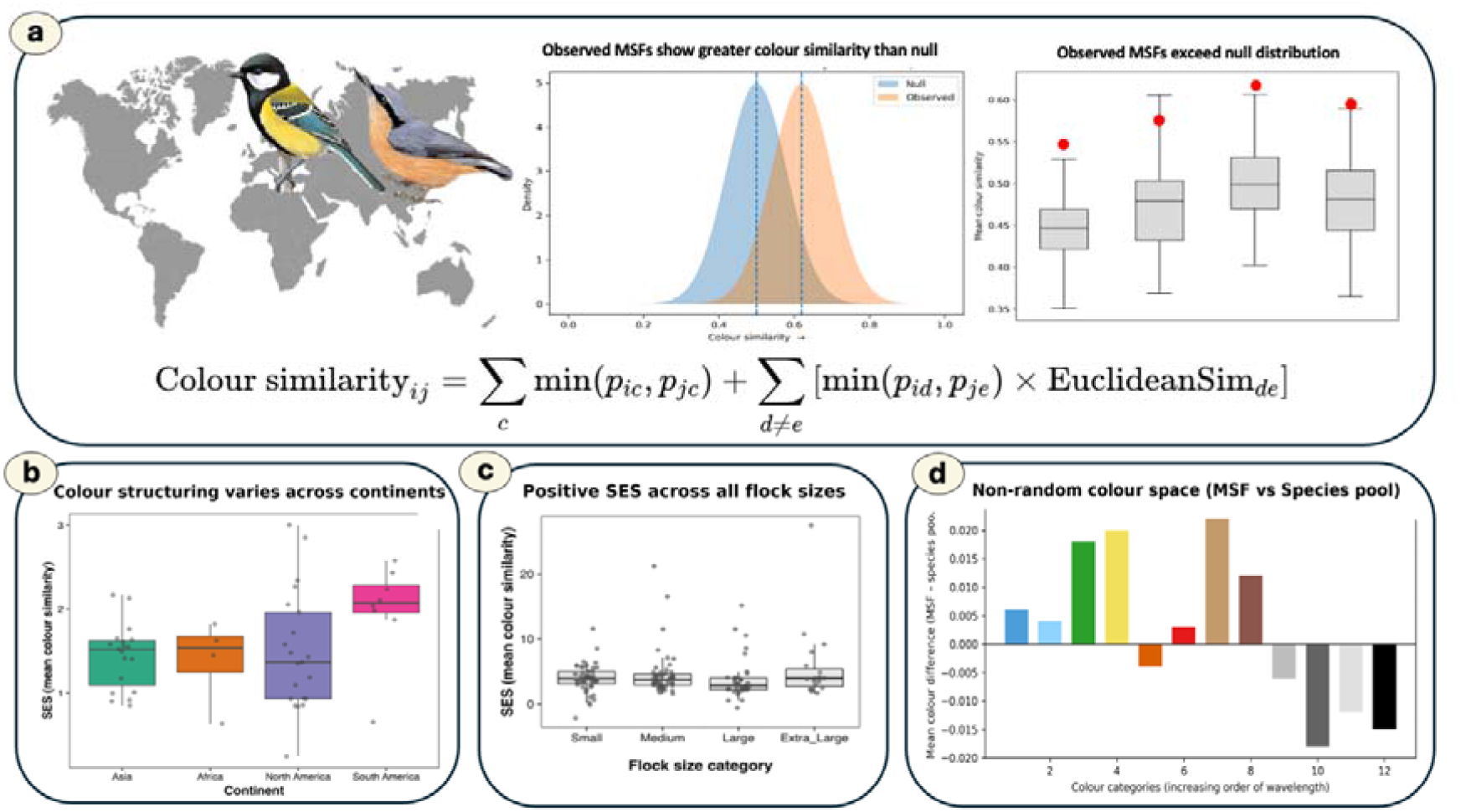
Global patterns of colour similarity in Mixed-Species Flocks (MSFs). This schematic summarises our global analysis of plumage colour structuring in 52 MSF datasets across four continents. Pairwise colour similarity was calculated using proportional overlap of colour categories (formula shown) to; (a) test whether observed flocks show greater mean colour similarity than null assemblages; (b) examine standardized effect sizes (SES) of colour structuring across flock-size categories and (c) explore the strength of structuring among continents. We also determined (d) whether MSF participants occupy a non-random region of colour space relative to regional species pools.

**Figure 2:**
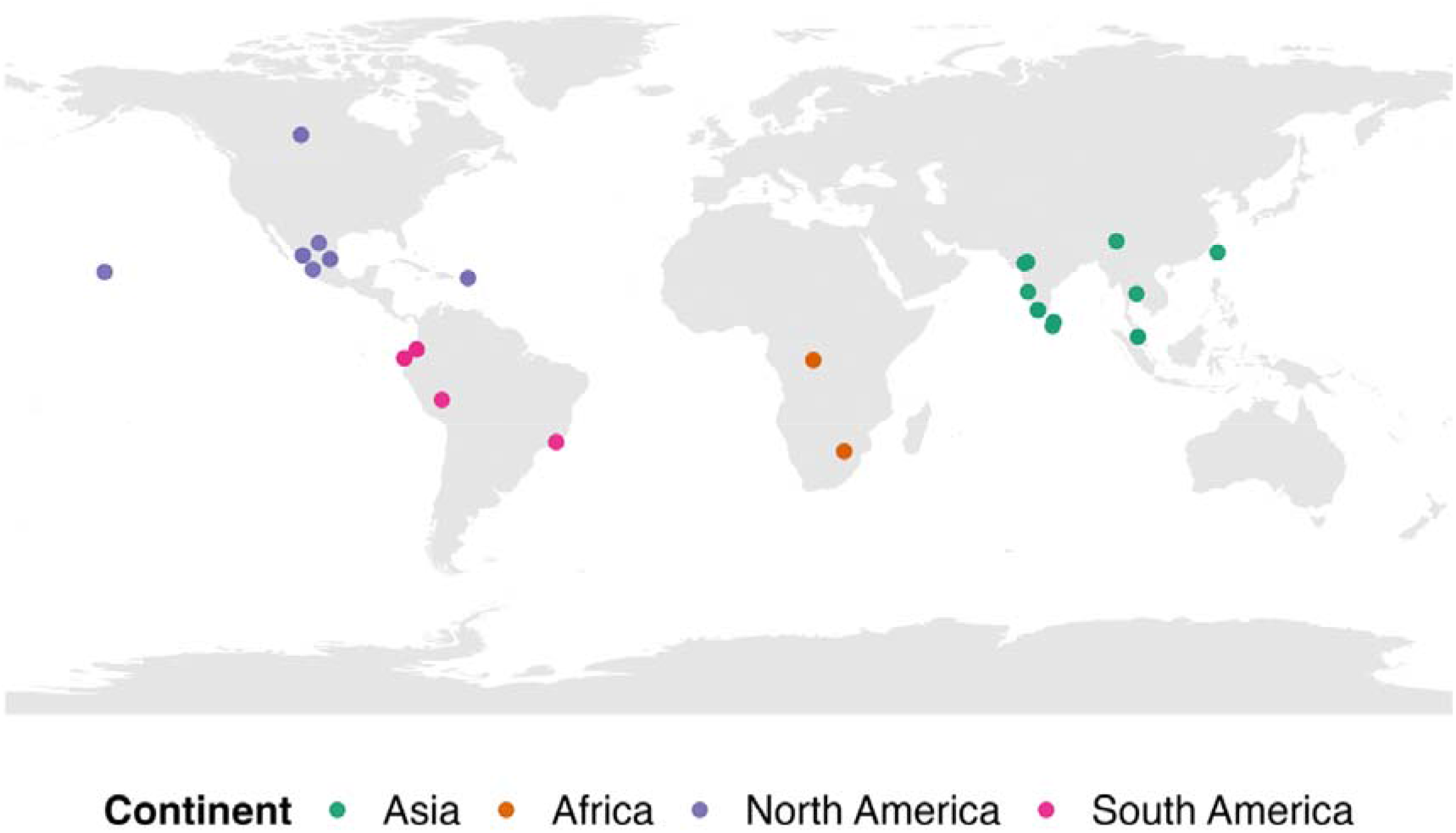
Global distribution of mixed-species flock (MSF) study sites. Geographic locations of the 52 MSF datasets compiled from 23 independent studies spanning four continents (Asia, Africa, North America, and South America). Each point represents a study site from which species-by-flock presence–absence matrices were derived. Together, these sites encompass a broad range of biogeographic regions and forest types, providing a globally distributed dataset for testing colour-based structuring in mixed-species bird flocks.

## Results

### Colour similarity in MSFs

Across all study sites and continents, MSFs consistently exhibited higher mean colour similarity than expected from randomly assembled flocks (Figure 3; Supplementary S2). In only 4 of 52 sites did observed values fall within the interquartile range (25–75% quantile) of the null distributions; these are Mexico_Coahuila (North America), Mexico_Western3 (North America), SouthAfrica_NylsvleyNatureReserve1 (Africa), and U.S._VirginIslands1 (North America) but observed values still lie above the mean of the null distributions. In all remaining sites, observed values exceeded the 75% quantile of the null expectation. This pattern was evident across Asia, Africa, North America, and South America, indicating that colour similarity among flock members is a widespread feature of mixed-species flocking rather than a region-specific phenomenon.

**Figure 3:**
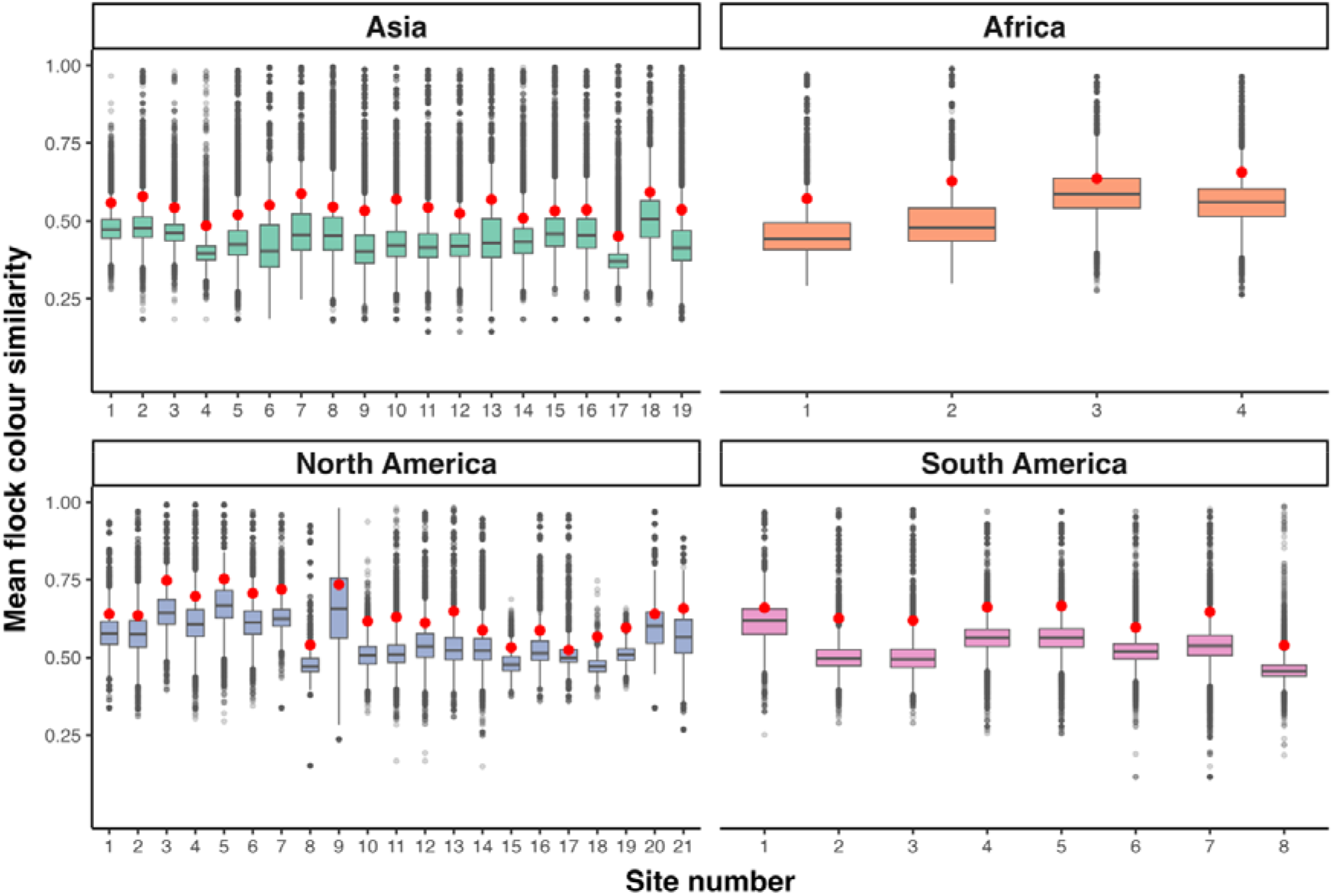
Colour similarity in mixed-species flocks at all sites. Site-wise comparisons of observed mean flock-level colour similarity (orange points) relative to null-model expectations (grey boxplots) across (a) Asia, (b) Africa, (c) North America, and (d) South America. Boxplots summarise the distribution of mean colour similarity across 1,000 null-model flocks per site. Observed values (orange points) indicate higher-than-expected colour similarity within mixed-species flocks at most sites.

This pattern is reinforced by examining the proportion of null-model flocks that exceeded observed colour similarity, which we determine by pseudo p, wherein values closer to zero indicate strong colour structuring (Fig. 4). Across all sites, 48 (out of 52) sites had a pseudo-p < 0.25, 24 sites had a pseudo-p < 0.1, and 12 sites had a pseudo-p < 0.05. These results indicate that observed colour similarity consistently lay in the upper tail of the null distribution across sites, providing strong evidence of non-random colour structuring in mixed-species flocks. Importantly, this metric directly quantifies the rarity of observed patterns under random assembly and provides site-specific evidence for colour-based structuring.

**Figure 4:**
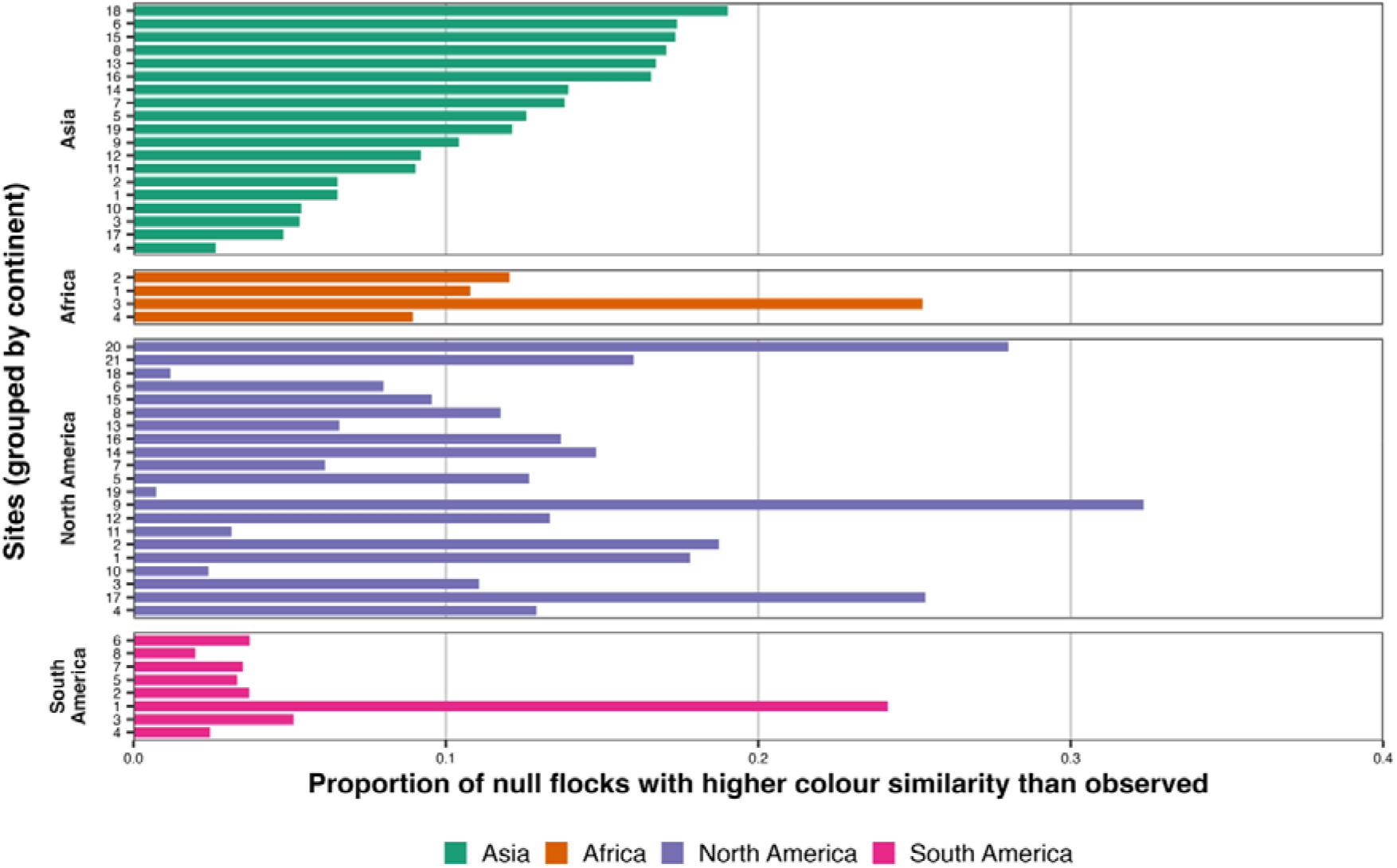
Frequency of null-model flocks exceeding observed colour similarity. Proportion of null-model flocks exhibiting higher mean colour similarity than observed mixed-species flocks, calculated separately for each site and grouped by continent. Values near zero indicate strong colour structuring, with observed flocks lying in the extreme upper tail of the null distribution. Across most sites, fewer than 20% of null flocks exceeded observed values, demonstrating that the observed colour similarity is unlikely to arise under random species assembly.

### Flock size & biogeography

SES values were consistently positive across all flock-size categories and continents (Fig. 5), indicating that observed flocks were more colour-similar than expected by chance regardless of species richness or geographic region. However, SES did not differ significantly among flock-size categories (ANOVA, F□,□□□ = 1.47, p = 0.226). Flock size explained only a small proportion of variance in SES (η² = 0.03). These results indicate that colour-based structuring operates similarly across flocks of different sizes, rather than strengthening or weakening with increasing richness.

**Figure 5:**
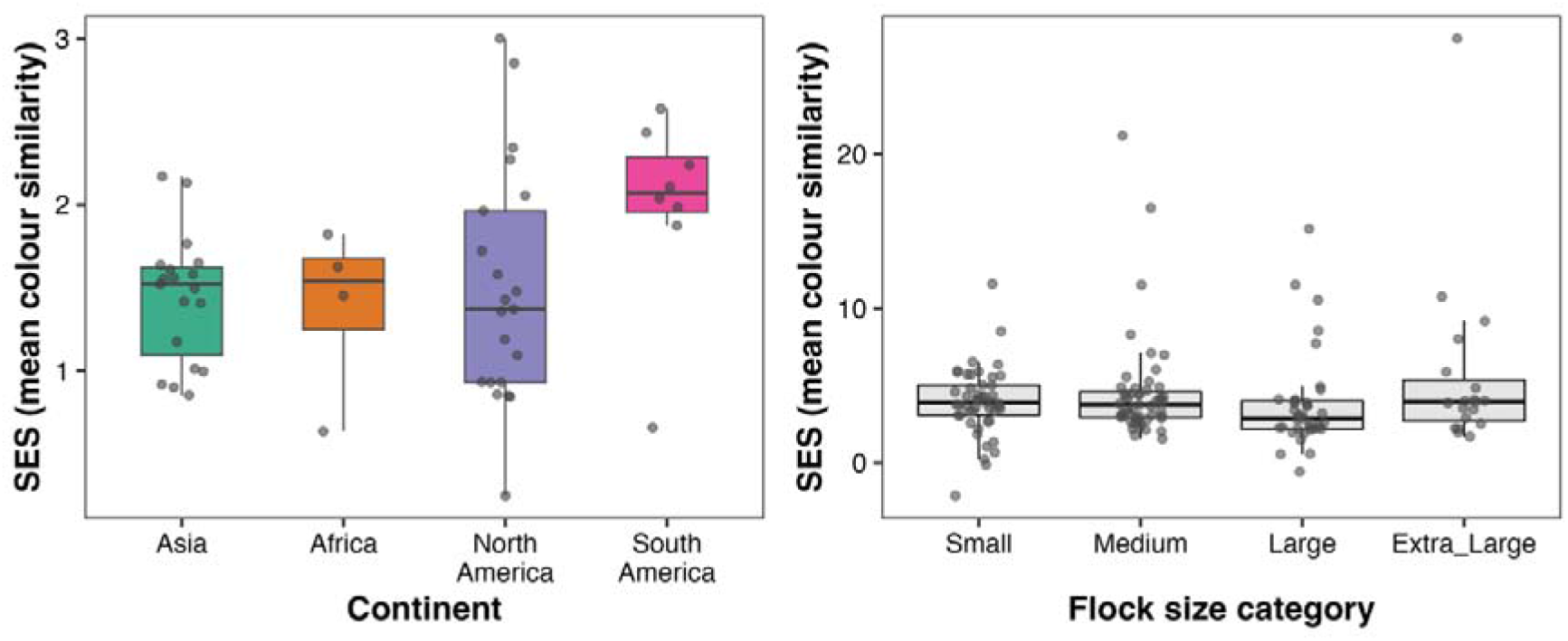
Flock size and biogeographic patterns in colour similarity of mixed-species bird flocks. Standardised effect sizes (SES) of mean within-flock colour similarity relative to null expectations across **(a)** continents and **(b)** flock size categories (small, medium, large, extra-large). Points represent site-level estimates; boxplots show medians and interquartile ranges. Positive SES values indicate greater colour similarity than expected under random assembly.

Similarly, colour structuring did not differ significantly among continents (ANOVA, F □,□□□ = 1.96, p = 0.133), although South America exhibited numerically higher SES values than other regions. Continent accounted for a modest proportion of variance in SES (η² = 0.11), but this effect was not statistically significant. Together, these findings suggest that colour-based structuring of MSFs is a geographically widespread phenomenon that does not vary systematically among major biogeographic regions.

Despite the absence of significant differences at the continental or flock-size level, site-level variation in SES indicates that the strength of colour structuring is not spatially uniform. Instead, the magnitude of structuring varies among local assemblages within continents (Supplementary information, Fig. S3)

### Colour in MSFs vs species pool

To assess whether MSF participants represent a colour-biased subset of the regional avifauna, we compared the proportional representation of 12 colour categories between MSF species and the local species pool across 52 sites spanning over 4 continents. Continent-level patterns were calculated as the mean proportional difference (MSF −species pool) across sites (Fig. 6, Supplementary Fig. S4). Positive mean proportional difference values indicate colours that are over-represented among MSF species, while negative mean proportional difference values indicate under-representation relative to locally available species.

**Figure 6:**
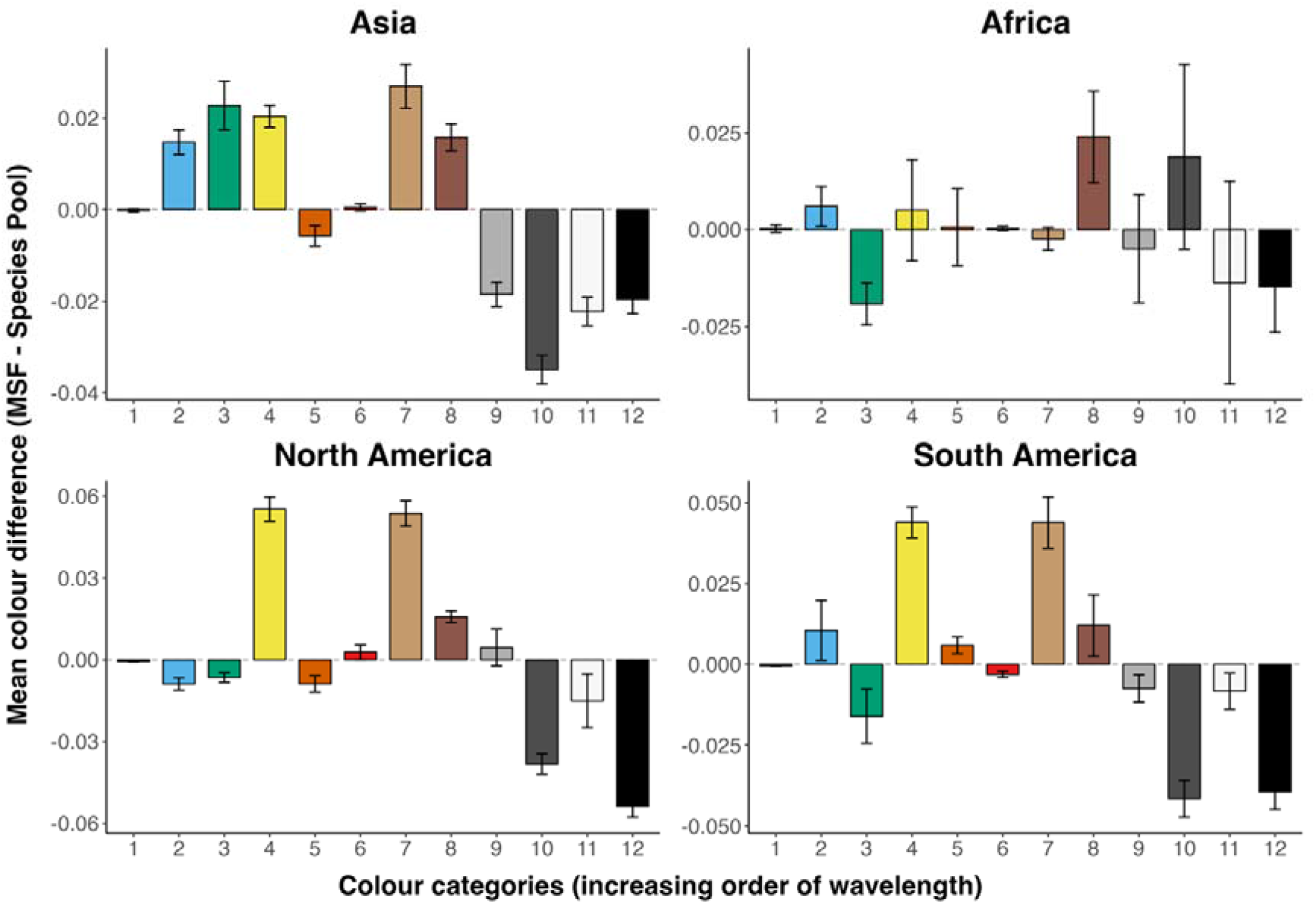
Colour composition of mixed-species flock participants relative to regional species pools. Bars show the mean difference in proportional representation of 12 colour categories between mixed-species flock (MSF) participants and the local species pool (MSF −species pool), averaged across sites within each continent. Positive values indicate colours that are over-represented among MSF species, while negative values indicate under-representation relative to locally available species. Colour categories are ordered by increasing wavelength.

Across all four continents, yellow and brown.dark were consistently over-represented, whereas black, white and grey.light were consistently under-represented. Grey.dark was under-represented and brown.light was over-represented in three of four regions, with Africa showing a partially distinct pattern. These continent-level trends represent average of patterns observed across individual study sites (Supplementary Fig. S4).

In Asian MSFs, chromatic colours, particularly brown.light (+0.027), green (+0.023), yellow (+0.020), brown.dark (+0.016), and blue (+0.015) were over-represented (Fig 6). In contrast, achromatic categories were consistently under-represented, particularly grey.dark (−0.035), white (−0.022), black (−0.020), and grey.light (−0.019). Rufous showed a slight reduction (−0.006), while red (+0.0005) and purple (−0.0002) exhibited negligible differences. Overall, Asian MSFs were characterized by increased representation of chromatic colours and reduced representation of achromatic colours (Fig 6).

North American MSFs exhibited the strongest over-represention of yellow (+0.055) and brown.light (+0.054) among all continents (Fig 6). Additionally, brown.dark (+0.016), grey.light (+0.004) and red (+0.003) were also over-represented. In contrast, strong under-representation was observed for black (−0.054) and grey.dark (−0.038), along with moderate reductions in white (−0.015), blue (−0.009), rufous (−0.009), and green (−0.007). Purple (−0.0005) showed minimal deviation. Thus, MSFs in North America were biased toward yellow–brown tones and depleted in achromatic colours (Fig 6).

South American MSFs (Fig 6) showed over-representation of yellow (+0.044) and brown.light (+0.044), followed by brown.dark (+0.012), blue (+0.011), and rufous (+0.006). In contrast, substantial under-representation was observed in grey.dark (−0.042) and black (−0.039). Smaller reductions were also evident for green (−0.016), white (−0.008), grey.light (−0.007), and red (−0.003). Purple (−0.0004) showed minimal deviation. Thus, MSFs in South America were also biased toward yellow–brown tones and depleted in achromatic colours (Fig 6).

African MSFs (Fig 6) displayed more moderate differences overall. Showed strong over-representation of brown.dark (+0.024) and grey.dark (+0.019), with smaller positive shifts in blue (+0.006) and yellow (+0.005). In contrast, colours such as green (−0.019), black (−0.015), white (−0.014), and grey.light (−0.005) were under-represented. Rufous (+0.0006), red (+0.0002), purple (+0.0002), brown.light (−0.002) showed a very small deviations. Compared to other continents, African MSFs showed weaker overall divergence from the species pool, with both chromatic and some darker achromatic colours contributing to MSF composition (Fig 6).

## Discussion

Plumage colour is a striking feature of birds and yet its role in structuring flocks is unexplored. We provide global evidence that plumage colour influences social community structure in MSFs. Across a global dataset spanning four continents, species participating in MSFs were found to be consistently more similar in colour than expected by chance, indicating that colour similarity is a widespread feature of flock composition. This pattern was remarkably consistent across flock sizes and geographic regions, suggesting that colour-based structuring is not restricted to particular ecological contexts but may reflect general visual processes influencing how birds associate in MSFs. Furthermore, MSF participants occupy a restricted region of colour space compared to the regional species pool, with yellow and brown colours consistently over-represented across continents. Together, these patterns suggest that colour filtering operates at multiple levels, shaping both the internal composition of flocks and the subset of species that participate in them.

### Colour similarity in MSFs

Across the 52 study sites spanning four continents, species participating in MSFs were consistently more similar in plumage colour than expected under random assembly. This pattern was observed across all regions included in our dataset and therefore does not appear to be restricted to any particular biogeographic context. These findings are broadly consistent with previous global analyses indicating that species associating within MSFs tend to be phenotypically similar. For example, the association strength among flocking species increases with similarity in traits such as body size and foraging behaviour across global MSF datasets^2^. In mixed species groups, phenotypic similarity is expected when protection from predators is gained through an increase in group size, known as supplementary benefits^2,5^. Our results extend this phenotypic similarity framework by showing that plumage colour represents an additional axis along which flock participants are structured. Taken together, these results indicate that MSF assembly may be influenced by multiple dimensions of phenotypic similarity, including morphology, behaviour, as well as visual traits.

Why might colour similarity matter in MSFs? Mixed-species flocks are highly dynamic groups that depend on rapid information transfer, coordinated movement, and mutual detection among heterospecific participants^5,8^. In visually complex environments such as forests, traits affecting detectability and signal recognition may therefore influence flock cohesion^13,25^. Species with broadly similar visual traits, including colour, may be more easily recognised as flock associates, and such similarity can potentially facilitate coordinated movement. Experimental studies in fish show that individuals preferentially associate with visually similar group members^26,27^. Whether similar mechanisms operate in mixed-species bird flocks remains largely untested. Similarity in plumage among flock members could also influence how flocks are perceived by predators. If individuals within a flock present a relatively uniform visual target, predator targeting efficiency may be reduced through mechanisms such as confusion or dilution effects^6,7,28^. Although our analyses do not directly test these mechanisms, the global consistency of colour similarity across MSFs suggests that visual traits may help species associate and move together within flocks.

Importantly, the observed pattern of colour similarity cannot be explained by uneven species commonness or variation in flock size, as both factors were explicitly controlled in the null-model framework. By maintaining proportional row (species presence) and column (flock composition) totals during our analysis, we preserved realistic species occurrence frequencies and flock richness distributions while randomising species associations^19,22^. As a result, the higher-than-expected colour similarity observed across sites likely reflects structured assembly rather than a statistical artefact. Overall, these analyses indicate that MSFs are more colour-similar than expected by chance and that colour similarity represents a widespread, non-random feature of MSF organisation at a global scale.

### Flock size and biogeographic context

Across all sites, colour structuring was consistently positive, and its magnitude did not vary significantly among flock-size categories or across continents. This suggests that colour-based assembly operates across a wide range of social contexts, from small MSFs containing only a few species to extra-large MSFs comprising more than 15 species. Colour similarity does not appear to weaken or strengthen with increasing flock richness but instead represents a consistent feature of MSF organisation. This is in contrast to previous work wherein phenotypic clumping among MSF participants was found to decrease as flock richness increases, suggesting that larger flocks may accommodate greater trait diversity^29^. In contrast, the consistency in colour similarity across flocks indicate that colour may represent a trait axis that remains structured even when other phenotypic traits diversify with increasing flock richness.

Even though species across continents differ greatly in their ecological conditions and evolutionary histories, the overall direction of colour structuring remained consistent. This broad geographic consistency suggests that the processes linking colour and flock participation are unlikely to be restricted to particular evolutionary radiations or habitat types, but may instead reflect more general principles of visual ecology in birds^13,25^. Variation among sites within continents, however, indicates that local ecological conditions likely modulate the strength of colour structuring. Habitat structure, light environment, predator communities, and local species composition may all influence how visual traits mediate interspecific associations^5,25^. The absence of strong inter-continental differences therefore does not imply uniformity across MSFs globally, but instead suggests that colour filtering is also influenced by factors at finer ecological scales. Together, these results indicate that while colour-based non-random assembly appears to be a widespread feature of MSFs, its strength may be shaped by local ecological conditions.

### The colours of MSF participants

Plumage colour in birds is shaped by a combination of ecological, evolutionary, and signalling processes, resulting in substantial variation in both conspicuousness and crypsis across species^30,31,32^. At the community level, colours of MSF participants did not represent a random subset of the regional avifauna. Instead, plumage colours of flocks were biased toward particular regions of colour space. Across all four continents, yellow and dark brown were consistently over-represented among MSF participants, whereas black, white, and light grey were consistently under-represented relative to the local species pool. Although colour representation in African MSFs showed somewhat weaker divergence from the species pool as compared to other continents, those MSFs were still non-random.

Interestingly, the colours that were most common among MSF participants differ from global patterns of colour prevalence across birds. Achromatic colours such as black, white, and grey are among the most widespread across bird species globally^9^, whereas our findings show that MSF participants under-represent black and white relative to the local species pool. Similarly, colours such as red and purple, which are often associated with sexual signalling^9^, showed minimal deviation between MSF participants and species pool. This likely reflects the fact that mixed-species flocks are not structured around mating interactions, but instead around anti-predator benefits. As a result, colours primarily involved in sexual signalling may play a limited role in determining participation in MSFs.

In contrast, MSF participants were disproportionately represented by yellow and brown across continents, which may be influenced by both pigmentation mechanisms and ecological functions relevant to flocking species. Yellow plumage in birds is typically produced by carotenoid pigments, which must be acquired through diet and are often associated with insect or plant-derived resources^33,34^. Because most MSF participants are predominantly insectivorous, carotenoid availability through diet may partly explain the frequent occurrence of yellow among MSF participants^35^. Brown plumage is usually produced by melanin pigments, which are widespread in birds and associated with feather durability as well as environmental adaptation^9,34^. These colours may also provide advantages in forest environments where most MSFs occur. Yellow and brown colours often blend well with foliage, bark, and leaf litter backgrounds, potentially enhancing camouflage and reducing detectability by predators across seasons^13,25^. Perceived conspicuousness further depends on differences in visual systems between birds and their predators^36^, suggesting that colours that appear important to conspecifics may remain less detectable to predators.

At a group level, flock participants that share broadly similar colour profiles increase visual uniformity within the flock, which can reduce predator targeting efficiency through confusion or dilution effects during attacks^6,28^. In contrast, strongly achromatic colours, such as black and white, are more conspicuous against natural backgrounds and adjacent colours^37^, compared to yellows and browns. Highly saturated colours such as red may also stand out within MSF, making individuals more visually distinct from other flock members and potentially more vulnerable to predators. Empirical studies have shown that conspicuous plumage can increase predation risk in some contexts^38^. However, this relationship is complex and context-dependent, varying with habitat structure and visual background^39^. Together, these findings suggest that both crypsis and conspicuousness operate along a continuum, and that the predominance of certain colour types in MSFs may reflect a balance between detectability, signalling, and the visual environment.

### Future directions

Although the patterns of colour similarity that we show here are remarkably consistent across MSFs globally, such analyses have inherent limitations. Species-specific colour proportions in this study were quantified from images of birds that were painted^9^ and therefore affected by how the images were created and analysed. They may not fully capture the range of intraspecific variation in plumage coloration. Also, in natural settings, plumage appearance can vary with lighting conditions, habitat structure, and viewing angle. Consequently, the degree of colour similarity estimated here is a coarse measure, and may miss some variation in how individuals are perceived in their natural environment^40^. Notably, the colour information used in this study does not incorporate UV, iridescence, and fluorescent colouration. However, such colours play an important role for sexual signalling and reproductive isolation^41^ and are designed to be conspicuous^42,43^, which is not expected to benefit members of MSFs. Finally, as with all global analyses, the dataset used in this study is geographically uneven, with most sites coming from Asia and North America, while Africa and South America are relatively under-represented. There are also major gaps within continents, with no representation from some regions of Europe, Australia, large parts of Central Asia and the Amazon basin.

Addressing some of the gaps in both the colour data and geographical spread would lead to more nuanced insights into the role of colour in flocks. Despite some limitations, our findings highlight how visual traits can influence community assembly in interspecific social systems. By integrating trait-based approaches with large-scale datasets, future studies may further clarify how visual signals interact with ecological and behavioural traits to structure mixed-species communities.

## Methods

### Colour similarity in MSFs

#### Dataset 1: MSF composition, flock categories and continents

We used a global dataset on species occurrences in mixed-species flocks (MSFs) compiled from 23 studies spanning four continents^2^ (Fig 2). The resulting compilation consists of 52 presence–absence matrices comprising 785 bird species, representing nearly 8% of all extant avian species. Each matrix represents a particular habitat type, elevational band, or spatially isolated patch. In each matrix, species are listed as rows and individual flocks as columns, with entries indicating presence (1) or absence (0) of each species in a given flock. Only species observed in at least one MSF within each study region were retained. For our analyses, we retained only species for which quantitative plumage colour data were available^9^, resulting in a final dataset of 682 species. The final dataset spans all four continents and includes the dominant taxonomic and ecological groups known to participate in MSFs, providing a robust global representation of MSF community structure.

To examine whether colour similarity varies with flock size, we categorised flocks based on species richness. Flocks containing 2–5 species were classified as small, 6–10 species as medium, 11–15 species as large, and flocks containing more than 15 species as extra-large. Flock size thresholds were based on richness quantiles across sites and adjusted to match natural breaks in the observed distributions. The number of flocks per site across continents and flock-size categories is summarised in Supplementary Table S1.

#### Dataset 2: Colour proportions

We used quantitative colour information for bird species, which estimated the proportional coverage of 12 discrete colour categories across the body surface, including plumage and exposed soft parts (bill, legs, and eyes) ^9^. Colour proportions were derived from RGB-based pixel classification applied to standardized bird plates from the *Handbook of the Birds of the World*, accessed via the *Birds of the World* platform^44^. The dataset comprises 22,325 images representing 10,618 species. Colours were classified into the following categories: blue, purple, red, yellow, green, rufous, light brown, dark brown, light grey, dark grey, black, and white. For each species, the proportion of pixels assigned to each colour category theoretically ranged from 0 to 1, although observed values spanned 0 to 0.97. These proportions provide a standardized, continuous representation of species-level colour composition suitable for comparative analyses across regions and assemblages.

#### Calculation of colour similarity

To quantify colour similarity among species participating in MSFs, we integrated the global data on MSF composition^2^, with the species-level plumage colour information^9^. For each pair of species within a flock, we calculated a continuous colour similarity score based on the overlap and similarity of colour in their plumage. Pairwise similarity was computed by first matching identical colour categories between species and weighting their overlap by the shared proportional coverage (Fig.1a). The remaining unmatched colours were then paired based on maximum pairwise-similarity between colours derived from a colour-similarity matrix (based on Euclidean similarity), ensuring that each colour component contributed only once to the final score. The overall colour similarity between two species was calculated as the sum of weighted similarities across all matched colour components, yielding a value bounded between zero (no similarity) and one (identical colour composition). Flock-level colour similarity was quantified by averaging pairwise similarity scores across all unique species pairs within a flock, yielding a mean colour similarity value per flock.

#### Pairwise colour similarity between species i and j

For each species, plumage colour was represented as the proportion of body surface covered by each colour category. Pairwise colour similarity between two species (i and j) was calculated by combining:

1. Exact matches of identical colours, and
2. Matches between different but maximally similar colours, determined by Euclidean similarity.

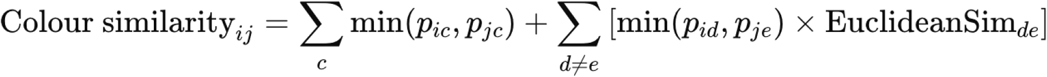

*where:*

- *“c” is the colour shared by both species i and j*.
- *“d” colour is present only in species i and “e” colour is present only in species j*.
- *“p” represent respective proportional plumage coverage of that colour category in respective species*.
- *the first term sums shared proportions of identical colour categories present in both species*.
- *the second term accounts for matches between different colour categories (d* ≠ *e), weighted by their Euclidean similarity*.

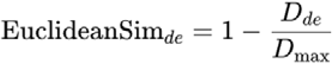

- D*_de_* is euclidean distance between colour categories *d* and *e*
- D_max_ is largest distance between colour categories that is theoretically possible (441.67 for RGB colour space)

In this equation, the minimum proportion of either identical or similar colours represents the proportion of that colour shared by the two birds. The similarity score was validated by matching the same species and species with completely different colours to obtain values of one and near zero. We also visually assessed pairs with different degrees of colour similarity to check whether the score performed as expected.

#### Null models and randomization algorithm

To test whether mixed-species flocks (MSFs) exhibit greater colour similarity than expected under random assembly, we implemented null models using the SIM8 algorithm in EcoSimR in R^19,20^. SIM8 is a proportional–proportional randomization algorithm in which both row totals (species occurrence frequencies) and column totals (flock sizes) are maintained proportional to their observed marginal totals. In species-by-flock matrices, this calculation ensures that the probabilities of species occurrence are conditional on both species-specific flocking tendencies and variation in flock richness among sites^21,22^. This choice is particularly appropriate for MSF datasets, where species differ markedly in their frequency and flocks vary substantially in richness. By conditioning on both row and column marginal totals we preserve realistic flock-size distributions and species-specific occurrence frequencies while randomizing co-occurrence structure. As a result, deviations in colour similarity reflect non-random association patterns rather than artefacts of uneven species abundance or variation in flock richness.

For each study site, we generated 1,000 randomized matrices from all flocking species at that site. For every null replicate, we calculated flock-level colour similarity using the same pairwise similarity metric applied to the observed data, thereby generating a null distribution of expected similarity values under random assembly. We then quantified the proportion of null replicates that produced colour similarity values greater than (or less than) the observed value, providing a site-specific and distribution-based estimate of the rarity of the observed pattern under the null model. Proportions above or below the observed value are considered as a pseudo ‘p’. Values of pseudo p closer to zero indicate stronger colour structuring, as they represent cases in which few null simulations produced higher similarity than observed, while larger values indicate weaker deviation from random assembly. Instead of using a strict significance threshold, these proportions are interpreted in a way similar to bootstrap support values in phylogenetic analyses, where results are viewed along a continuum rather than as simply significant or non-significant. In this way, the values reflect the strength of deviation from the null expectation and allow comparison across sites.

#### Standardized effect size

To quantify the magnitude and direction of deviation of observed flock-level colour similarity from null expectations, we calculated standardized effect sizes (SES). For each site, SES was calculated as the difference between the mean observed colour similarity and the mean colour similarity obtained from the null model, divided by the standard deviation of the null distribution:

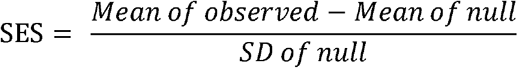

SES can vary from positive to negative values where “positive” values indicate that observed flocks are more colour-similar than expected by chance, and “negative” values indicate lower-than-expected similarity. SES values provide a standardised measure of effect size that allows comparison of the strength of colour structuring across sites that differ in flock size distributions and species composition. Rather than using SES as a strict hypothesis-testing statistic, we interpreted SES values as continuous measures of deviation from null expectations. Values exceeding ±1.96, corresponding to the outer 95% of the null distribution^23^, were used as a reference to identify particularly strong deviations, but emphasis was placed on the direction, magnitude, and consistency of SES patterns across flock-size categories and biogeographic regions using ANOVA. SES values were therefore used to compare the relative strength of colour structuring across sites, flock sizes, and continents in subsequent analyses.

### Colour in MSFs vs species pool

#### Dataset 3

Mixed-species flock (MSF) participants are widely recognised as representing a non-random subset of the local avian community, shaped by ecological, behavioural, and functional constraints on flock membership^1,3,24^. To assess whether this selectivity extends to plumage colour composition, we compiled regional species pools for each study site. Species occurrence data were obtained from the Global Biodiversity Information Facility (GBIF), using geographic coordinates corresponding to each study site^2^. For each of the 52 sites, we extracted all bird species recorded within the corresponding geographic area to represent the local species pool. This approach allowed us to characterise the full set of species potentially available to participate in MSFs at each site.

Species from both the MSF datasets and the regional species pools were matched to species-level plumage colour composition data^9^, ensuring consistent colour information across all comparisons. Using these data, we quantified and compared the average proportional representation of each of the 12 human-visible colour categories between MSF participants and the regional species pool, calculated separately for each site and summarised across continents.

#### Average colour proportions

To quantify differences in colour composition between mixed-species flock (MSF) participants and the regional species pool, we calculated average plumage colour proportions at the site and continental scales. For each species, colour composition was represented as the proportional coverage of 12 human-visible colour categories (blue, purple, red, yellow, green, rufous, light brown, dark brown, light gray, dark gray, black, and white)^9^.

At each study site, we first calculated the mean proportional representation of each colour category across all MSF-participating species. Species-level colour proportions were averaged without weighting by flock frequency to ensure that each species contributed equally to the site-level estimate, independent of its abundance or frequency of occurrence in flocks. In parallel, we calculated the mean proportional representation of each colour category across all species in the corresponding regional species pool, using the same averaging procedure. For each site and colour category, we then calculated the difference in mean colour proportion between MSF participants and the regional species pool (MSF mean −species pool mean). This yielded a site-level estimate of whether particular colour categories were over- or under-represented among MSF participants relative to locally present species.

To facilitate broader geographic comparisons, site-level colour differences were subsequently averaged within continents (Asia, Africa, North America, and South America). All calculations were conducted using identical procedures for MSF participants and species pool assemblages to ensure comparability. All analyses were carried out using R Version 4.1.1.

## Acknowledgements

We thank Umesh Srinivasan for his continued guidance. We also thank Md. Abdus Shakur for assistance with the GBIF platform, and Geetika Aggarwal for her constant companionship and discussions. K.A. acknowledges support from the Prime Minister’s Research Fellowship during her PhD at the Centre for Ecological Sciences, Indian Institute of Science.

## Author Contributions

Conceptualization: K.A., M.T., and K.S.

Methodology: K.A., I.S., M.T., and K.S.

Investigation: K.A.

Visualization: K.A., I.S., M.T., and K.S.

Supervision: M.T., and K.S.

Writing—original draft: K.A.

Writing—review & editing: K.A., M.T., and K.S.

