## Supplementary material for "Plumage colour structures global patterns of mixed-species bird flocks"

**Supplementary Table S1.** Number of mixed-species flocks per site across continents and flock-size categories. Flock sizes were classified based on species richness (small: 2–5; medium: 6–10; large: 11–15; extra-large: >15 species)

| Continent | Site no. | Sites | Total MSFs | Total Species | Flock size categories | | | | GBIF reference for Species pool datasets |
|---|---|---|---|---|---|---|---|---|---|
| | | | | | Small | Medium | Large | Extra large | |
| Asia | 1 | India_Anaimalaihills1 | 28 | 32 | 1 | 12 | 13 | 2 | GBIF.org (07 October 2024) GBIF Occurrence Download https://doi.org/10.15468/dl.v7fjyz |
| Asia | 2 | India_Anaimalaihills2 | 28 | 33 | 5 | 8 | 15 | 0 | GBIF.org (07 October 2024) GBIF Occurrence Download https://doi.org/10.15468/dl.v7fjyz |
| Asia | 3 | India_Anaimalaihills3 | 38 | 38 | 3 | 10 | 22 | 3 | GBIF.org (07 October 2024) GBIF Occurrence Download https://doi.org/10.15468/dl.v7fjyz |
| Asia | 4 | India_Anaimalaihills4 | 30 | 57 | 1 | 5 | 12 | 12 | GBIF.org (07 October 2024) GBIF Occurrence Download https://doi.org/10.15468/dl.v7fjyz |
| Asia | 5 | India_Anshi | 188 | 54 | 66 | 63 | 40 | 19 | GBIF.org (8 October 2024) GBIF Occurrence Download https://doi.org/10.15468/dl.jyhncm |
| Asia | 6 | India_GujaratPurva | 24 | 24 | 21 | 3 | 0 | 0 | GBIF.org (8 October 2024) GBIF Occurrence Download https://doi.org/10.15468/dl.46thza |
| Asia | 7 | India_GujaratRatanmahal | 29 | 28 | 23 | 5 | 1 | 0 | GBIF.org (30 May 2025) GBIF Occurrence Download https://doi.org/10.15468/dl.jstq4s |
| Asia | 8 | India_Namdapha | 95 | 53 | 44 | 49 | 2 | 0 | GBIF.org (8 October 2024) GBIF Occurrence Download https://doi.org/10.15468/dl.87hfyj |
| Asia | 9 | India_Parambikulam1 | 116 | 47 | 67 | 41 | 7 | 1 | GBIF.org (8 October 2024) GBIF Occurrence Download https://doi.org/10.15468/dl.jz6hqu |
| Asia | 10 | India_Parambikulam2 | 88 | 50 | 34 | 45 | 6 | 3 | GBIF.org (8 October 2024) GBIF Occurrence Download https://doi.org/10.15468/dl.jz6hqu |
| Asia | 11 | Malaysia_Fraser'sHill1 | 23 | 33 | 9 | 7 | 6 | 1 | GBIF.org (8 January 2026) GBIF Occurrence Download https://doi.org/10.15468/dl.gsckwc |
| Asia | 12 | Malaysia_Fraser'sHill2 | 23 | 35 | 5 | 11 | 6 | 1 | GBIF.org (8 January 2026) GBIF Occurrence Download https://doi.org/10.15468/dl.gsckwc |
| Asia | 13 | Malaysia_Fraser'sHill3 | 29 | 31 | 20 | 6 | 3 | 0 | GBIF.org (8 January 2026) GBIF Occurrence Download https://doi.org/10.15468/dl.gsckwc |
| Asia | 14 | SriLanka_KnucklesRange1 | 26 | 42 | 7 | 12 | 5 | 0 | GBIF.org (12 January 2026) GBIF Occurrence Download https://doi.org/10.15468/dl.xmgsvf |
| Asia | 15 | SriLanka_KnucklesRange2 | 27 | 36 | 8 | 17 | 2 | 0 | GBIF.org (12 January 2026) GBIF Occurrence Download https://doi.org/10.15468/dl.xmgsvf |

| | | | | | | | | | |
|---|---|---|---|---|---|---|---|---|---|
| Asia | 16 | SriLanka_KnucklesRange3 | 38 | 27 | 20 | 17 | 1 | 0 | GBIF.org (12 January 2026) GBIF Occurrence Download https://doi.org/10.15468/dl.xmgsvf |
| Asia | 17 | SriLanka_Sinharaja | 152 | 40 | 19 | 70 | 50 | 13 | GBIF.org (4 January 2026) GBIF Occurrence Download https://doi.org/10.15468/dl.agckf9 |
| Asia | 18 | Taiwan_Fushanexperimentalforest | 120 | 29 | 84 | 34 | 2 | 0 | GBIF.org (12 January 2026) GBIF Occurrence Download https://doi.org/10.15468/dl.c467ag |
| Asia | 19 | Thailand_KhaoyaiNP | 72 | 56 | 41 | 25 | 6 | 0 | GBIF.org (12 January 2026) GBIF Occurrence Download https://doi.org/10.15468/dl.a6sy3d |
| Africa | 1 | Congo_SalongaNationalPark1 | 18 | 25 | 10 | 8 | 0 | 0 | GBIF.org (2 January 2026) GBIF Occurrence Download https://doi.org/10.15468/dl.fpcd4p |
| Africa | 2 | Congo_SalongaNationalPark2 | 21 | 26 | 18 | 3 | 0 | 0 | GBIF.org (2 January 2026) GBIF Occurrence Download https://doi.org/10.15468/dl.fpcd4p |
| Africa | 3 | SouthAfrica_NylsvleyNatureReserve1 | 47 | 30 | 34 | 13 | 0 | 0 | GBIF.org (9 January 2026) GBIF Occurrence Download https://doi.org/10.15468/dl.rhebd7 |
| Africa | 4 | SouthAfrica_NylsvleyNatureReserve2 | 87 | 35 | 54 | 33 | 0 | 0 | GBIF.org (9 January 2026) GBIF Occurrence Download https://doi.org/10.15468/dl.rhebd7 |
| North America | 1 | Canada_Saskatchewan2 | 13 | 16 | 10 | 3 | 0 | 0 | GBIF.org (2 January 2026) GBIF Occurrence Download https://doi.org/10.15468/dl.vb2muh |
| North America | 2 | Canada_Saskatchewan3 | 14 | 31 | 7 | 7 | 0 | 0 | GBIF.org (2 January 2026) GBIF Occurrence Download https://doi.org/10.15468/dl.vb2muh |
| North America | 3 | Canada_Saskatchewan4 | 24 | 25 | 17 | 6 | 1 | 0 | GBIF.org (2 January 2026) GBIF Occurrence Download https://doi.org/10.15468/dl.vb2muh |
| North America | 4 | Canada_Saskatchewan5 | 41 | 39 | 24 | 15 | 2 | 0 | GBIF.org (2 January 2026) GBIF Occurrence Download https://doi.org/10.15468/dl.vb2muh |
| North America | 5 | Canada_Saskatchewan6 | 55 | 29 | 44 | 11 | 0 | 0 | GBIF.org (2 January 2026) GBIF Occurrence Download https://doi.org/10.15468/dl.vb2muh |
| North America | 6 | Canada_Saskatchewan7 | 16 | 30 | 9 | 4 | 3 | 0 | GBIF.org (2 January 2026) GBIF Occurrence Download https://doi.org/10.15468/dl.vb2muh |
| North America | 7 | Canada_Saskatchewan8 | 8 | 23 | 4 | 3 | 1 | 0 | GBIF.org (2 January 2026) GBIF Occurrence Download https://doi.org/10.15468/dl.vb2muh |
| North America | 8 | Hawaii_Hakalauwildliferefuge | 28 | 7 | 26 | 2 | 0 | 0 | GBIF.org (8 January 2026) GBIF Occurrence Download https://doi.org/10.15468/dl.54d6jg |
| North America | 9 | Mexico_Coahuila | 64 | 35 | 63 | 1 | 0 | 0 | GBIF.org (13 January 2026) GBIF Occurrence Download https://doi.org/10.15468/dl.bpwxzg |
| North America | 10 | Mexico_ElCielobiospherereserve1 | 9 | 31 | 1 | 5 | 2 | 1 | GBIF.org (05 October 2023) GBIF Occurrence Download https://doi.org/10.15468/dl.fmbwsw |
| North America | 11 | Mexico_ElCielobiospherereserve2 | 43 | 39 | 7 | 22 | 13 | 1 | GBIF.org (05 October 2023) GBIF Occurrence Download https://doi.org/10.15468/dl.fmbwsw |

| | | | | | | | | | |
|---|---|---|---|---|---|---|---|---|---|
| North America | 12 | Mexico_ElCielobiosphereres erve3 | 22 | 27 | 9 | 12 | 1 | 0 | GBIF.org (05 October 2023) GBIF Occurrence Download https://doi.org/10.15468/dl.fmbwsw |
| North America | 13 | Mexico_ElCielobiosphereres erve4 | 15 | 25 | 7 | 5 | 3 | 0 | GBIF.org (05 October 2023) GBIF Occurrence Download https://doi.org/10.15468/dl.fmbwsw |
| North America | 14 | Mexico_Jalisco | 55 | 20 | 20 | 25 | 8 | 2 | GBIF.org (12 January 2026) GBIF Occurrence Download https://doi.org/10.15468/dl.ajkk62 |
| North America | 15 | Mexico_Western1 | 5 | 21 | 0 | 3 | 1 | 1 | GBIF.org (8 January 2026) GBIF Occurrence Download https://doi.org/10.15468/dl.uf2wte |
| North America | 16 | Mexico_Western2 | 7 | 11 | 6 | 1 | 0 | 0 | GBIF.org (8 January 2026) GBIF Occurrence Download https://doi.org/10.15468/dl.uf2wte |
| North America | 17 | Mexico_Western3 | 10 | 9 | 5 | 5 | 0 | 0 | GBIF.org (8 January 2026) GBIF Occurrence Download https://doi.org/10.15468/dl.uf2wte |
| North America | 18 | Mexico_Western4 | 4 | 21 | 0 | 2 | 1 | 1 | GBIF.org (8 January 2026) GBIF Occurrence Download https://doi.org/10.15468/dl.uf2wte |
| North America | 19 | Mexico_Western5 | 11 | 37 | 0 | 0 | 3 | 8 | GBIF.org (8 January 2026) GBIF Occurrence Download https://doi.org/10.15468/dl.uf2wte |
| North America | 20 | U.S._VirginIslands1 | 31 | 11 | 29 | 2 | 0 | 0 | GBIF.org (12 January 2026) GBIF Occurrence Download https://doi.org/10.15468/dl.sbuqt2 |
| North America | 21 | U.S._VirginIslands2 | 6 | 10 | 5 | 1 | 0 | 0 | GBIF.org (12 January 2026) GBIF Occurrence Download https://doi.org/10.15468/dl.6z7pvd |
| South America | 1 | Brazil_Teresopolis | 8 | 19 | 4 | 4 | 0 | 0 | GBIF.org (05 October 2023) GBIF Occurrence Download https://doi.org/10.15468/dl.rb9ghy |
| South America | 2 | Ecuador_GuanderaBiological Reserve1 | 30 | 28 | 8 | 13 | 9 | 0 | GBIF.org (12 January 2026) GBIF Occurrence Download https://doi.org/10.15468/dl.e8wcgm |
| South America | 3 | Ecuador_GuanderaBiological Reserve2 | 17 | 23 | 8 | 5 | 3 | 1 | GBIF.org (12 January 2026) GBIF Occurrence Download https://doi.org/10.15468/dl.e8wcgm |
| South America | 4 | Ecuador_MachalillaNationalP ark1 | 112 | 28 | 23 | 86 | 3 | 0 | GBIF.org (04 October 2023) GBIF Occurrence Download https://doi.org/10.15468/dl.kzgpne |
| South America | 5 | Ecuador_MachalillaNationalP ark2 | 110 | 24 | 34 | 69 | 7 | 0 | GBIF.org (04 October 2023) GBIF Occurrence Download https://doi.org/10.15468/dl.kzgpne |
| South America | 6 | Ecuador_MachalillaNationalP ark3 | 97 | 45 | 5 | 43 | 46 | 3 | GBIF.org (04 October 2023) GBIF Occurrence Download https://doi.org/10.15468/dl.kzgpne |
| South America | 7 | Ecuador_MachalillaNationalP ark4 | 115 | 38 | 25 | 69 | 19 | 2 | GBIF.org (04 October 2023) GBIF Occurrence Download https://doi.org/10.15468/dl.kzgpne |
| South America | 8 | Peru_Cochacashu | 32 | 94 | 2 | 2 | 4 | 24 | GBIF.org (8 January 2026) GBIF Occurrence Download https://doi.org/10.15468/dl.5emb4f |

**Supplementary Figure S2. Observed versus null distributions of mean colour similarity in mixed-species flocks (MSFs)**

Density distributions of mean flock-level colour similarity for observed mixed-species flocks (orange) and null-model flocks (blue) across four continents: (a) Asia, (b) Africa, (c) North America, and (d) South America. Null distributions were generated from 1,000 randomised species assemblages per site that preserved flock size and species occurrence frequencies. Dashed vertical lines indicate mean values of the observed (orange) and null (blue) distributions for each site. Across continents, observed flocks consistently show higher colour similarity than expected under random assembly, indicating non-random colour structuring of flock composition.

#### Asia

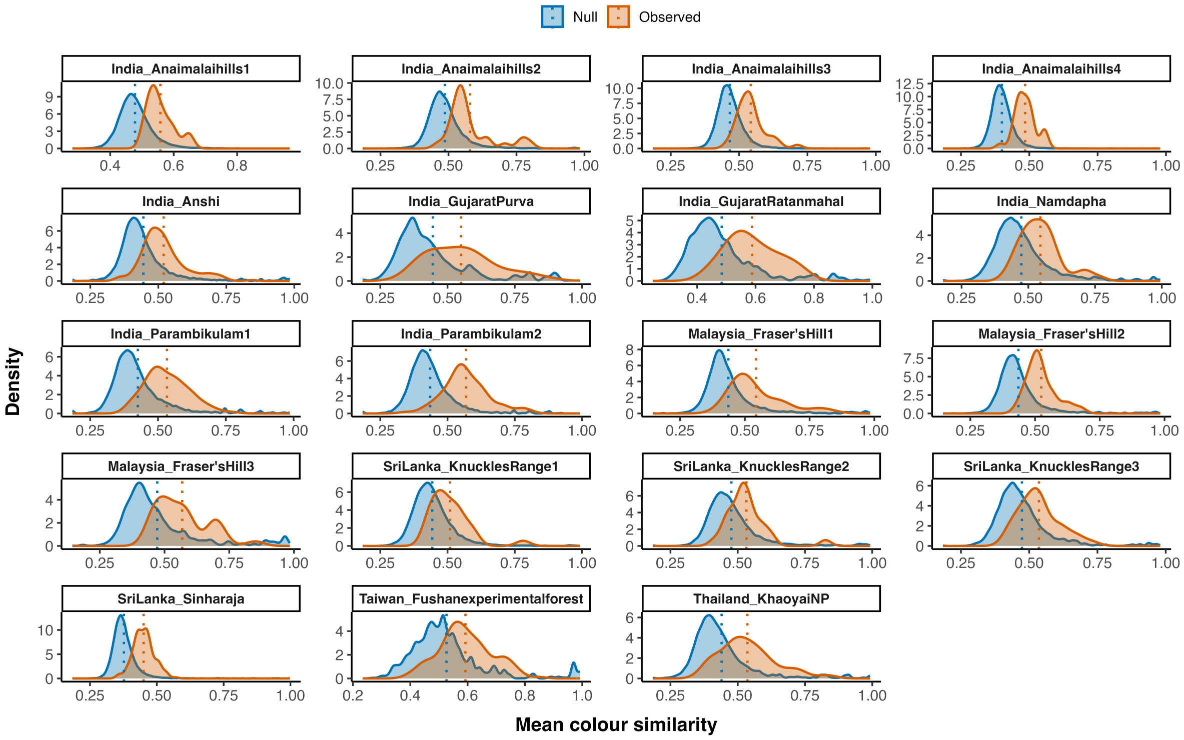

Null
Observed
India_Anaimalaihills1
India_Anaimalaihills2
India_Anaimalaihills3
India_Anaimalaihills4
India_Anshi
India_GujaratPurva
India_GujaratRatanmahal
India_Namdapha
India_Parambikulam1
India_Parambikulam2
Malaysia_Fraser'sHill1
Malaysia_Fraser'sHill2
Malaysia_Fraser'sHill3
SriLanka_KnucklesRange1
SriLanka_KnucklesRange2
SriLanka_KnucklesRange3
SriLanka_Sinharaja
Taiwan_Fushanexperimentalforest
Thailand_KhaoyaiNP
Density
Mean colour similarity


#### Africa

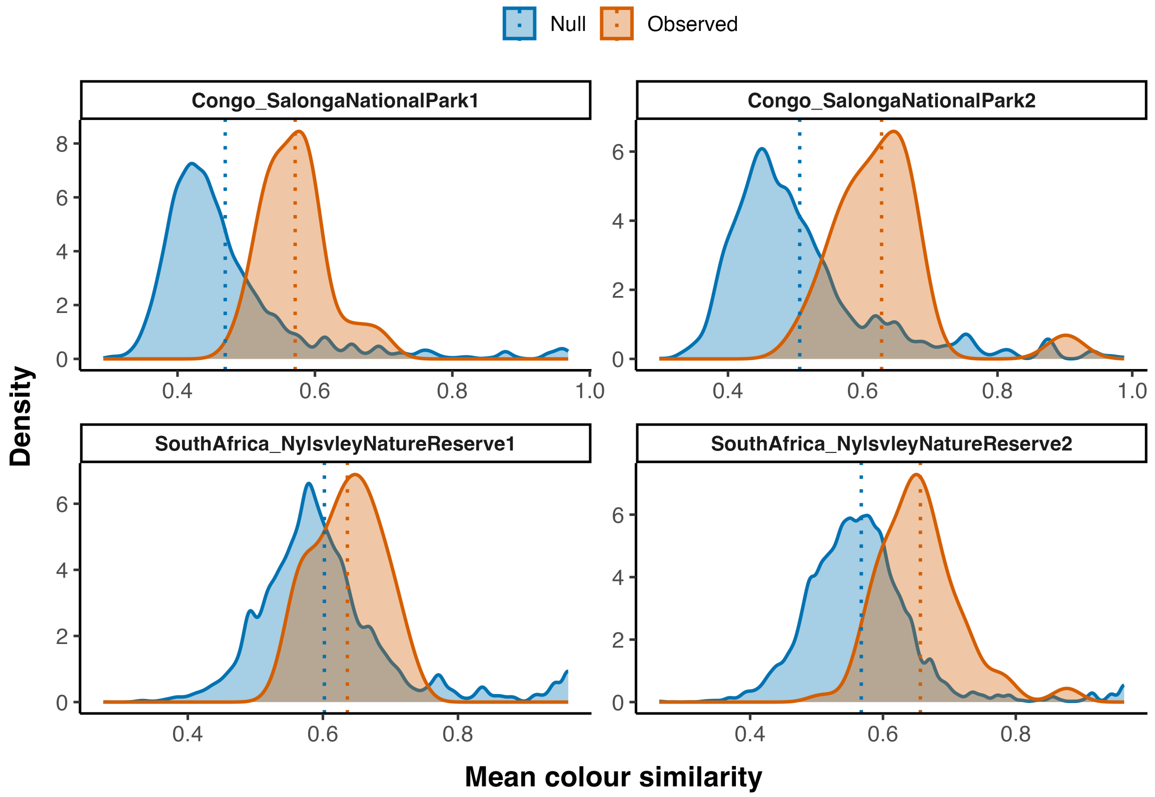

Null
Observed
Congo_SalongaNationalPark1
Congo_SalongaNationalPark2
SouthAfrica_NylsvleyNatureReserve1
SouthAfrica_NylsvleyNatureReserve2
Density
Mean colour similarity

### North America

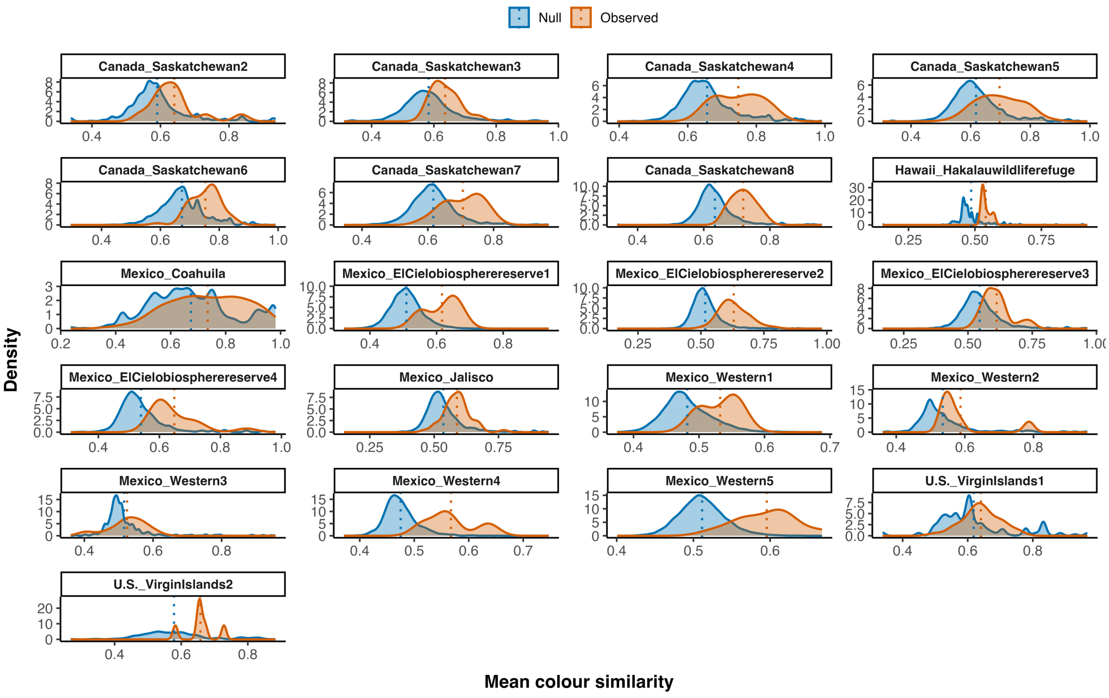


### South America

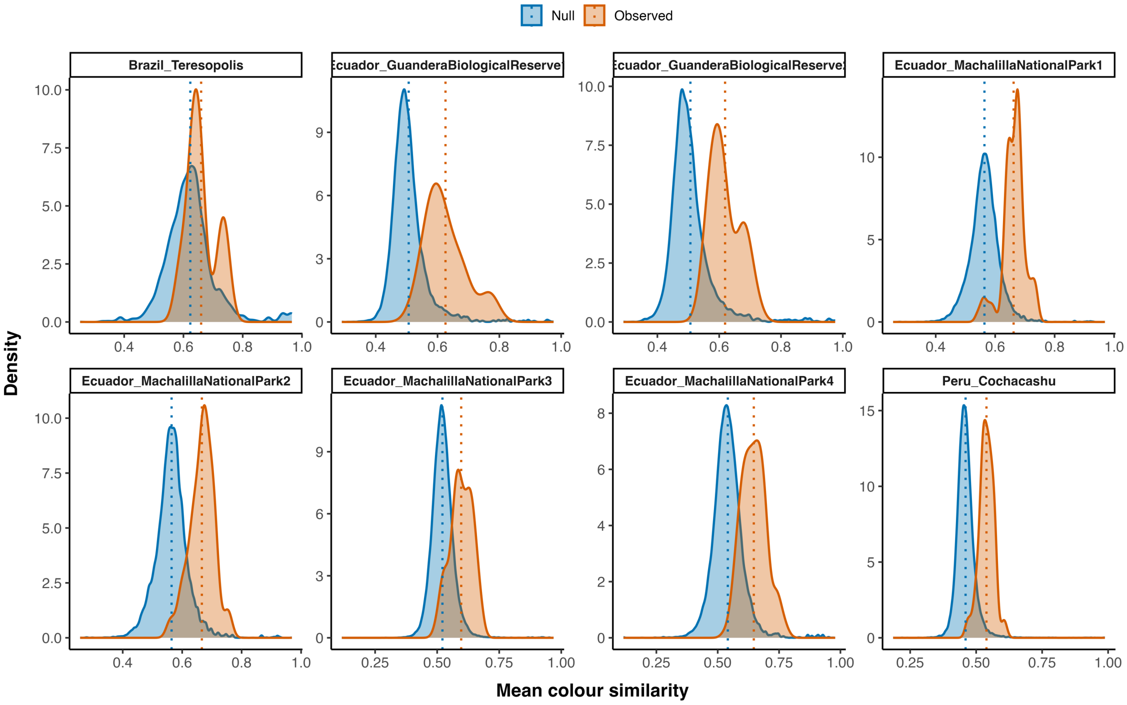

**Supplementary Figure S3. Site-level colour structuring of mixed-species flocks across flock sizes and continents**

Each point represents the standardised effect size (SES) of mean colour similarity for a single site and flock-size category. Colours denote flock-size classes (small, medium, large, extra-large), and sites are grouped by continent (Asia, Africa, North America, South America). The vertical dashed line at SES = 0 indicates no deviation from null expectations.

This figure illustrates substantial heterogeneity in the strength of colour structuring among sites within continents and across flock sizes, while reinforcing the general tendency for observed flocks to be more colour-similar than expected under random assembly. This site-level variation complements the continental and flock-size summaries shown in Fig. 5.

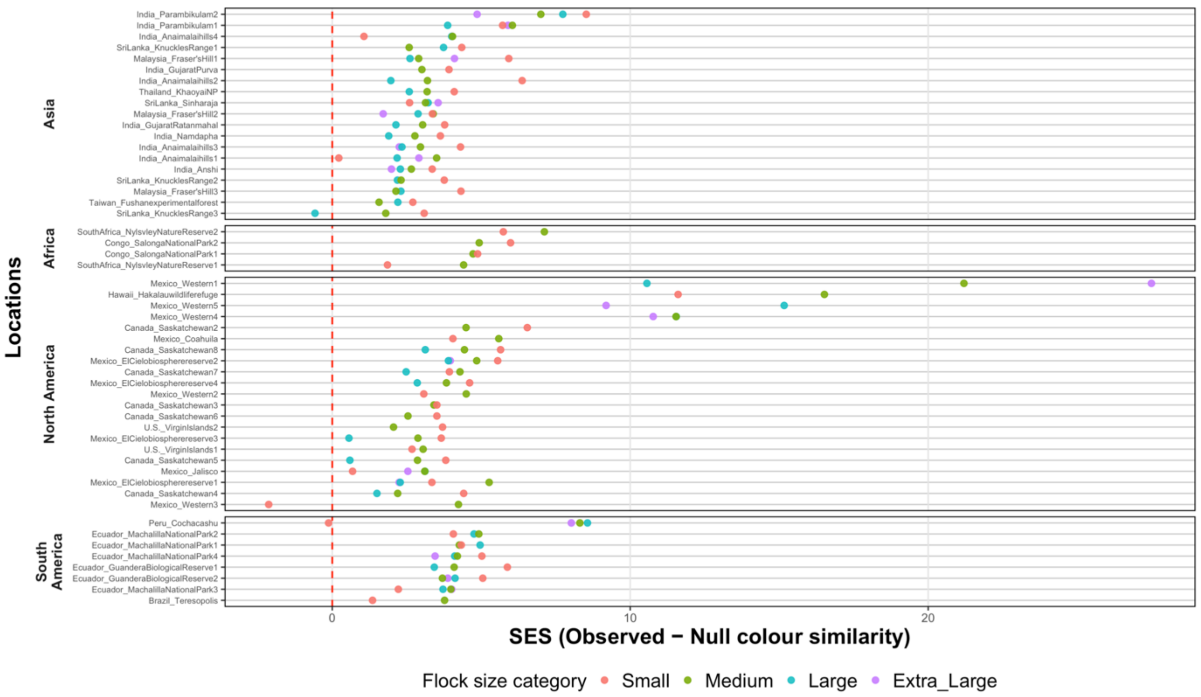

**Supplementary Figure S4. Site-level colour composition differences between mixed-species flock participants and regional species pools.**

Bars show the difference in mean proportional representation of each colour category between MSF participants and the local species pool (MSF − species pool) calculated separately for each study site. These plots illustrate within-continent heterogeneity underlying the continent-level patterns presented in Fig. 6.

### Colour differences by site - Asia

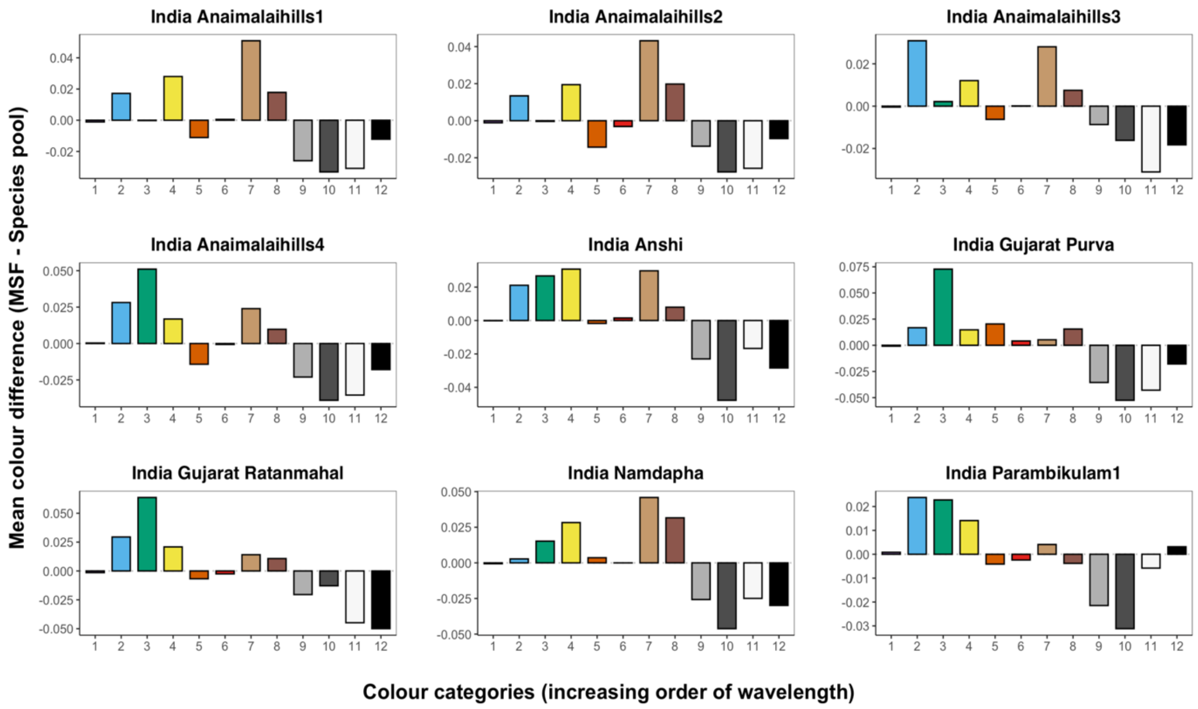


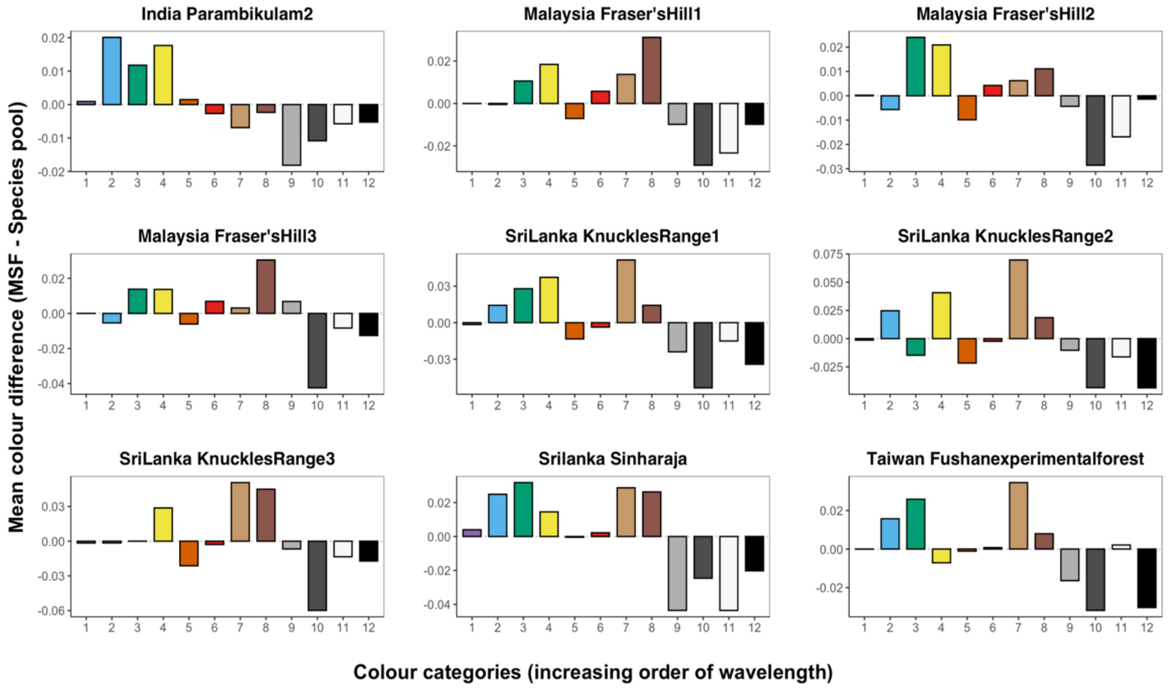


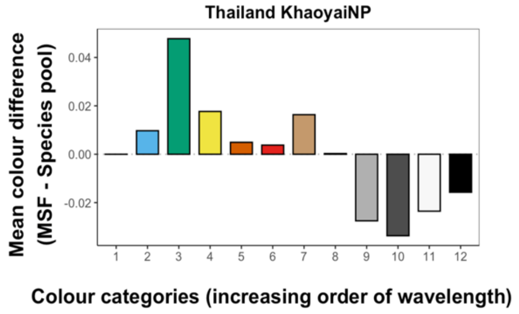

### Colour differences by site – Africa

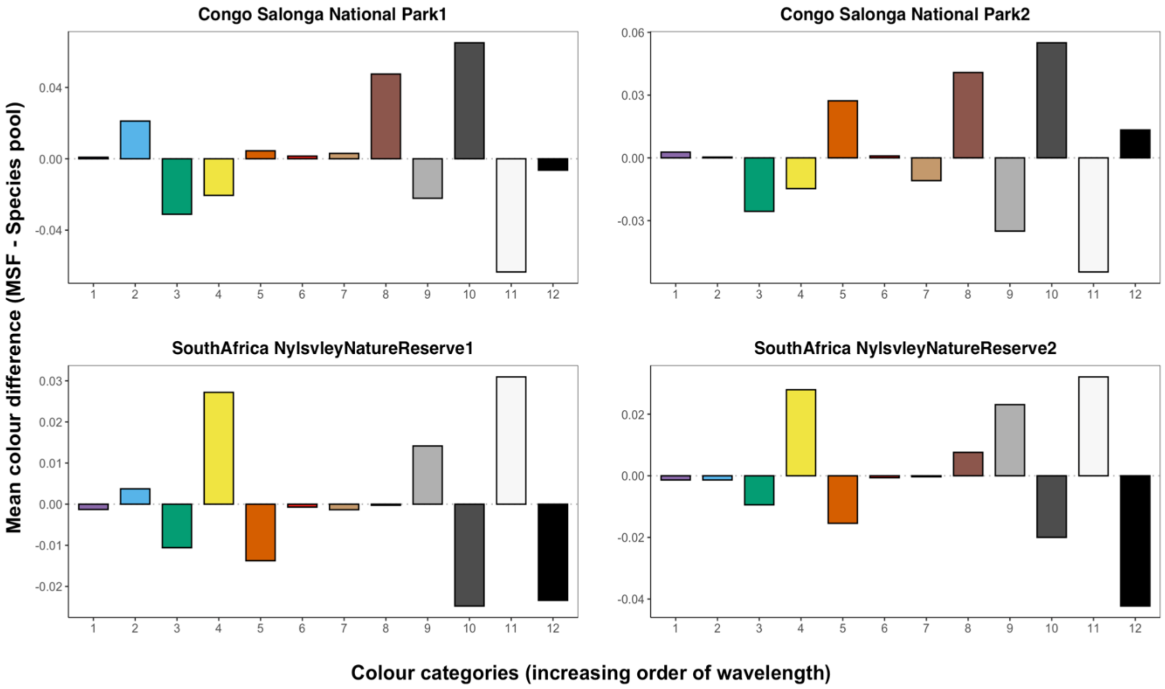

### Colour differences by site – North America

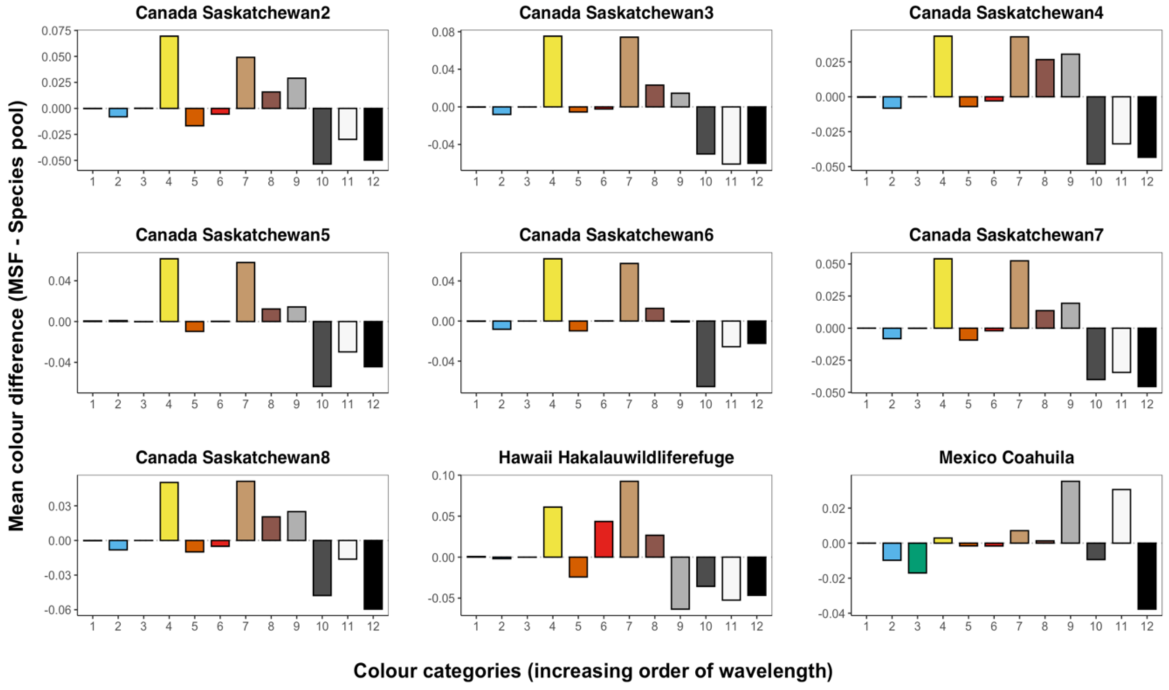


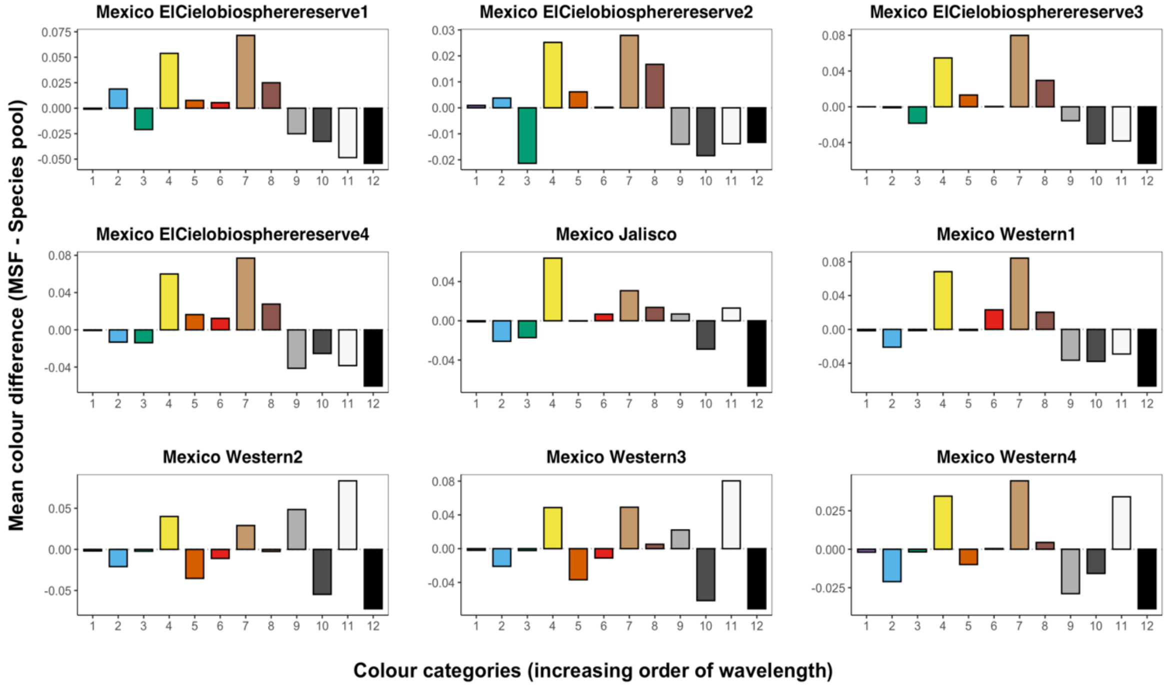


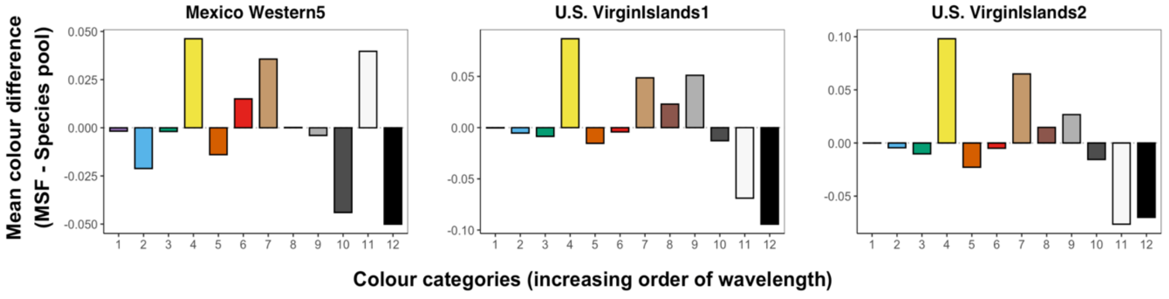

### Colour differences by site – South America

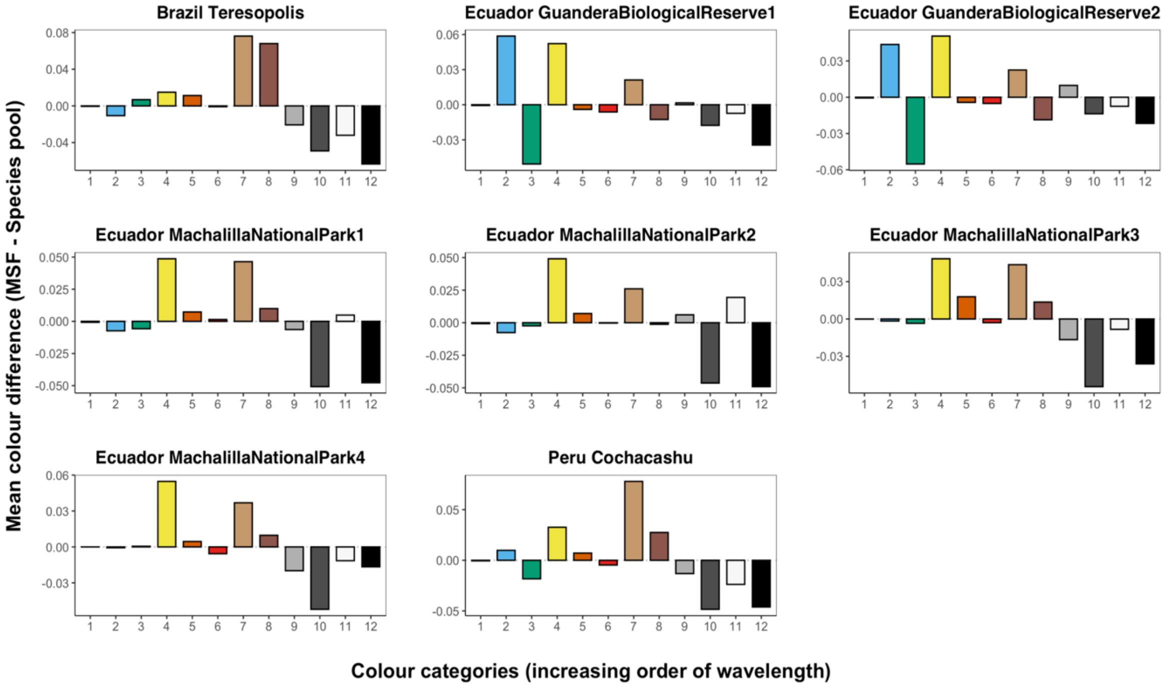
